# An integrative approach to investigate natural variation in the accumulation of aliphatic glucosinolates in *Arabidopsis thaliana*

**DOI:** 10.1101/838227

**Authors:** Suraj Sharma, Ovidiu Popa, Stanislav Kopriva, Oliver Ebenhoeh

## Abstract

Glucosinolates are a fascinating class of specialised metabolites found in the plants of *Brassicacea* family. The variation in glucosinolate composition across different Arabidopsis ecotypes could be a result of allelic compositions at different biosynthetic loci. The contribution of methylthioalkylmalate synthase (MAM) genes to diversity of glucosinolate profiles across different Arabidopsis ecotypes has been confirmed by genetic analyses. Different MAM isoforms utilise different chain-elongated substrates for glucosinolate biosynthesis causing thus a variation in chain lengths across different Arabidopsis ecotypes. To further investigate the relationship between the genotype and the associated metabolic phenotype, we studied the diversity of genes and enzymes of glucosinolate biosynthesis. Using Shannon entropy as a measure we revealed that several genes of the pathway show a clear derivation from the expected behaviour, either accumulating non-synonymous SNPs or showing signs of purifying selection. We found that the genotype-phenotype relationship is much more complicated than inferred from the diversity of MAM synthases. We conclude therefore, that the ON/OFF feature of key QTLs is not enough to elucidate the diversity of glucosinolates across different *Arabidopsis thaliana* ecotypes and that glucosinolate profiles are determined also through the polymorphic residues along the coding regions of multiple metabolic genes.

## Introduction

Plant secondary metabolism produces a huge variation in structures and molecules with a plethora of functions for the plants but also for human nutrition and health (Owen, Patron, Huang, & Osbourn, 2017). One class of such secondary metabolites are glucosinolates in the Brassicaceae. Glucosinolates (GSLs) are important for the plants as a defence against herbivores, fungi, and other pathogens (Halkier & Gershenzon, 2006). They are also determinants of taste and flavour of cruciferous vegetables and responsible for their positive health properties (Traka & Mithen, 2009). More than 140 different GSL structures have been described, with a great variation not only between species, but also among ecotypes of the same species (Agerbirk et al., 2015; Clarke, 2010). GSLs are synthesised from amino acids and are divided into three classes: aliphatic GSLs, derived from alanine, leucine, isoleucine, valine and methionine (Met), indolic GSLs synthesised from tryptophan, and aromatic GSLs from phenylalanine (Phe) (Halkier & Gershenzon, 2006; Sønderby, Geu-Flores, & Halkier, 2010). All GSLs possess the same core structure, which comprises a glucose residue linked to a (Z)-N-hydroximic sulfate ester via a sulphur atom (Fahey, Zalcmann, & Talalay, 2001). GSL biosynthesis consists of three independent steps: (i) chain elongation of selected precursor amino acids (only Met and Phe), (ii) formation of the core GSL structure, and (iii) secondary modifications of the amino acid side chain. The diversity of GSLs derives from the side-chain elongation and secondary modifications.

GSLs are best known for their function in interaction between plants and herbivores. Upon tissue damage, the GSLs stored in the vacuoles come in contact with the enzyme myrosinase, which cleaves the thio-glucoside bond. The resulting aglycones are unstable and form volatile isothiocyanates or nitriles (Halkier & Gershenzon, 2006). Depending upon the herbivore, the volatiles of specific GSLs can act as feeding deterrents or stimulants (Buskov, Serra, Rosa, Sørensen, & Sørensen, 2002; Gabrys & Tjallingii, 2002; Lazzeri, Curto, Leoni, & Dallavalle, 2004; Mewis, Ulrich, & Schnitzler, 2002; Miles, Campo, & Renwick, 2005; Noret et al., 2005). Therefore, there is often an overlap between quantitative trait loci (QTL) for GSL accumulation and insect resistance (Kroymann, Donnerhacke, Schnabelrauch, & Mitchell-Olds, 2003). A possible outcome of this heterogeneous natural selection on GSLs is the quick evolution of new compounds or new patterns of compound accumulation (Daxenbichler et al., 1991; Rodman, 1980). New GSLs may increase resistance to herbivores that have become adapted to existing defences, whereas new patterns of GSL accumulation may provide a unique complement of defences by slowing down the counter-adaptation of herbivores.

The GSL defence system is one of the few systems where systematic data assessing between and within species variation at both phenotypic and causal genetic level is available (Halkier & Gershenzon, 2006; Sønderby et al., 2010). Natural variation within or between species is regulated by a complex network of genes and associated polymorphisms (Fisher, 1930; Kliebenstein, Kroymann, et al., 2001; Lynch, Walsh, & others, 1998). These variations, however, complicate our understanding of how certain genes behave in context of a species as we often study a single genotype. Thus, understanding a metabolic pathway requires studies involving more than one ecotype. For example, methionine derived aliphatic GSLs differ in length of the side chain caused by the elongation of the amino acid, as well as by further modifications, e.g. oxidation of the sulfur atom (Halkier & Gershenzon, 2006; Sønderby et al., 2010). In the model plant *Arabidopsis thaliana*, aliphatic GSLs of six different chain-lengths, referred to as 3C to 8C GSLs, but with different side chains are found. The diversity in length of aliphatic GSLs can be explained by the variation in the iterative chain-elongation cycles, each adding one methylene group to the Met and/or elongated Met molecule (Halkier & Gershenzon, 2006). The QTL responsible for determining the chain-elongation of GSLs is GS-ELONG (Magrath et al., 1994). GS-ELONG is highly variable across different Arabidopsis ecotypes, with common large indel polymorphism (Kroymann et al., 2003). The gene underlying the GS-ELONG QTL is methylthioalkylmalate synthase (MAM3), encoding the key enzyme of the chain elongation cycle (Kroymann et al., 2001). However, the GS-ELONG locus harbours two more genes, isoforms of MAM3 called MAM1 and MAM2. MAM3 is present in all Arabidopsis ecotypes, and some ecotypes possess both additional genes whereas others possess either MAM1 or MAM2, with some, such as *Landsberg erecta* (*Ler*), having a truncated (non-functional) MAM1 in addition to MAM2 (Kroymann et al., 2003). While the presence of functional MAM genes has been described as responsible for the variation in aliphatic GSL side chain length, very little is known about contribution of other genetic variation, mainly single nucleotide polymorphism (SNP), at this locus.

In this paper, we investigate the link between the diversity of GSL enzyme-coding genes and their GSL profiles exhibited across 72 different *A. thaliana* ecotypes. The selection of 72 ecotypes is based on the availability of information about the gene sequences and the patterns of accumulation of aliphatic glucosinolates. Importantly, the experiments for determining the aliphatic glucosinolate levels have been performed under identical conditions (Chan, Rowe, & Kliebenstein, 2010; Kliebenstein, Kroymann, et al., 2001). It can therefore be assumed that the environment was identical (up to experimental precision) for all ecotypes. This study presents an effort to quantify the impact of the diversity of MAM gene sequence on GSL variation, rather than the on/off nature of the GS ELONG QTLs. The genetic sequence coding for an enzyme determines the kinetic properties of the corresponding enzyme. For example, polymorphisms in the active sites, in principle, can change the substrate specificity of the respective enzyme. Thus, we investigate the level of polymorphisms in the coding region of the GSL enzymes to study the impact on the diversity of aliphatic GSLs.

## Results

### Distribution of MAM genes across Arabidopsis thaliana ecotypes

The genetic basis of chain-length distribution of aliphatic GSLs became evident with the identification of the GS-ELONG QTL in Arabidopsis and *Brassica napus* (Magrath et al., 1994). The locus was mapped in Arabidopsis by using a cross between two ecotypes, *Columbia (Col)* and *Landsberg erecta (Ler)*, where the major glucosinolates are *4C* and *3C*, respectively (Kroymann et al., 2001). The underlying candidates *MAM1* and *MAM3* genes are two adjacent sequences with high similarity to genes encoding isopropylmalate synthase that catalyses the condensation of chain elongation in leucine biosynthesis. Later, a third MAM-like gene, *MAM2*, was identified at the same locus as *MAM1* (Kroymann et al., 2003). While *MAM3* is ubiquitous, most Arabidopsis ecotypes examined possessed functional copies of either *MAM1* or *MAM2* genes. A functional *MAM1* gene has been correlated with the accumulation of 4C GSLs, whereas a functional *MAM2* has been linked to 3C GSLs. To gain more insights on the impact of MAM synthases on chain lengths distribution of aliphatic GSL, we analysed the similarity of the annotated *MAM1* gene across 72 Arabidopsis ecotypes taken from 1001 genome project database (Jorge et al., 2016). These ecotypes were selected based on the availability of gene sequences and the associated aliphatic GSL profiles. Details on the 72 ecotypes are given in the Supplementary Table 1. The annotated *MAM1* sequences from the 72 ecotypes were compared to each other for diversity. Figure 1 shows a mid-point rooted phylogenetic tree showing the evolutionary relationship between the ecotypes based on the similarities and differences in the coding region of the MAM1/MAM2 sequences. Based on maximum likelihood estimation (Guindon et al., 2010), the tree shows two main branches. While 53 out of the 72 ecotypes clustering in the blue branch indeed possess high similarity to the coding region of *MAM1* gene, 19 ecotypes in the red branch possess genes more like the *MAM2* gene. Thus, we assume that the ecotypes composed in blue and red branches possess *MAM1* and *MAM2* genes, respectively.

**Figure 1:**
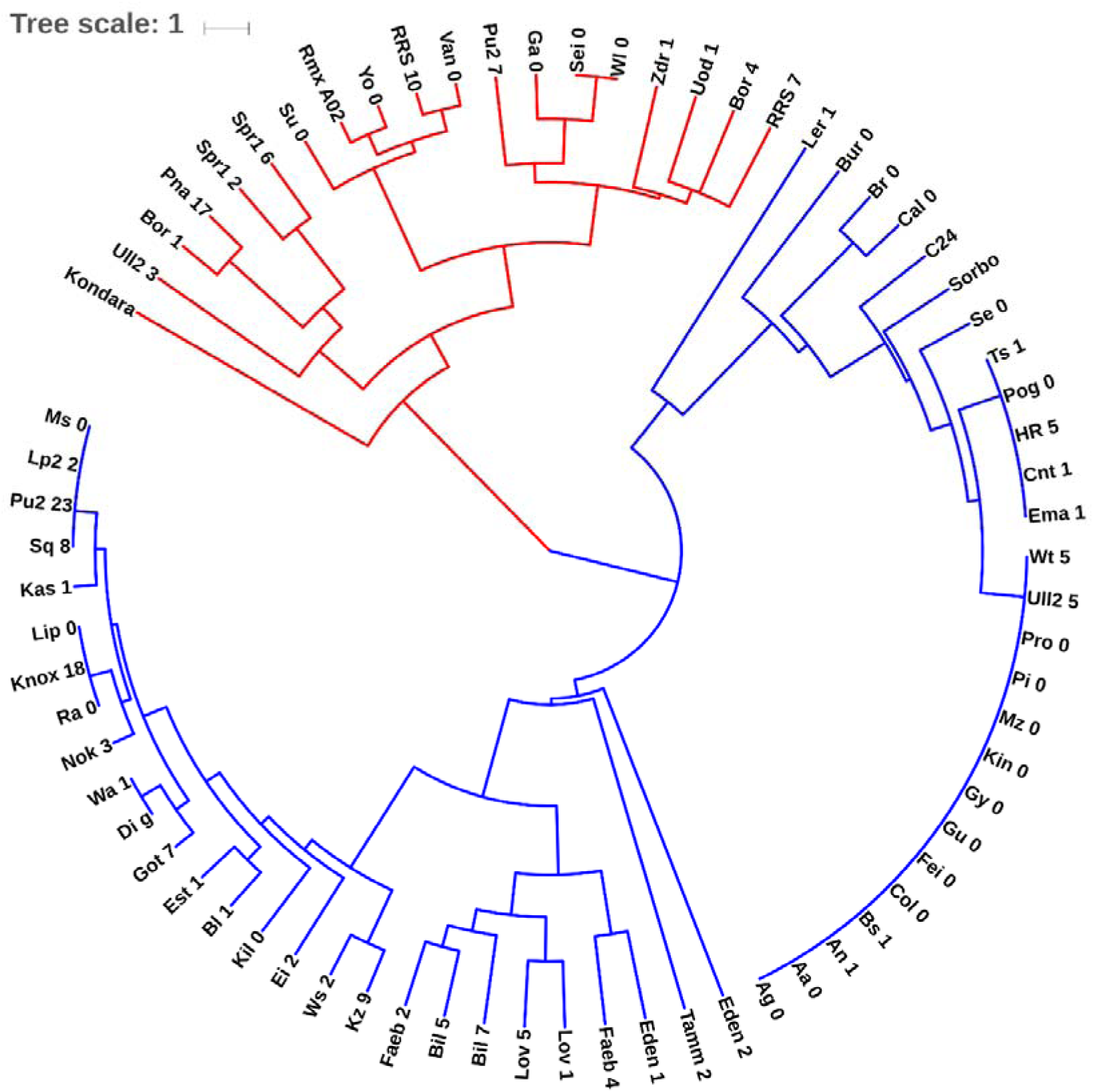
Mid-point rooted phylogenetic tree showing the evolutionary relationship among coding regions of annotated MAM1 gene of 72 Arabidopsis thaliana ecotypes. The red branch represents ecotypes showing higher similarity to the MAM2 sequence. The scale bar is substitutions per position.

### The metabolic genotypes and associated phenotypes

We define a metabolic genotype (*G*_*i*_) as the gene sequence of enzymes of glucosinolate biosynthesis in ecotype *i*, whereas the metabolic phenotype (*P*_*i*_) corresponds to the composition of aliphatic GSLs in the ecotype *i*. To gain a deeper understanding of how different metabolic genotypes and their associated phenotypes are linked, we analysed the genotypic and phenotypic distances. The genotypic distances between the genotypes were calculated as Hamming distance (Hamming, 1950) 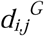, which is the number of positions at which the corresponding nucleotide/amino-acid characters are different between gene sequences *G*_*i*_ and *G*_*j*_ of equal length. The phenotypic distance 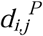 was calculated as Euclidean distance, 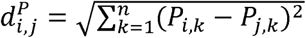 between two phenotypes *P*_*i,k*_ = (*P*_*i*,1_,*P*_*i*,2_,…, *P*_*i,n*_) and *P*_*j,k*_ = (*P*_*j,1*_,*P*_*j,2*_,…, *P*_*j,n*_) in an *n*-dimensional space. In this study, n=6, which corresponds to the total number of chain-elongated aliphatic GSLs found in *A. thaliana* (Figure 2). Thus, we can quantify differences between all pairs of ecotypes based on their metabolic genotype *G*_*i*_ (*i* = 1, …,72) and phenotype *P*_*i*_ (*i* = 1,…, 72). Figure 3 showcases the summary of the analysis, where the genotypic distances are plotted against the phenotypic distances. Intuitively, one would assume that similar genotypes shall exhibit similar phenotypes, and *vice-versa*. However, Figure 3 clearly shows that several ecotypes show low genotypic distance (i.e. they are genotypically similar) but exhibit high phenotypic distance (variation in GSL profiles). Also, there exist genotypically diverse ecotypes that exhibit very similar GSL profiles. Thus, investigating the factors affecting variations in the phenotypes of such ecotypes will provide a clearer understanding of how distinct patterns of GSL accumulation emerge out of genetic differences. Moreover, investigation of the localisation of polymorphic residues in the GSL biosynthesis enzymes will provide a better understanding of the link between metabolic genotype and the associated metabolic phenotypes.

**Figure 2:**
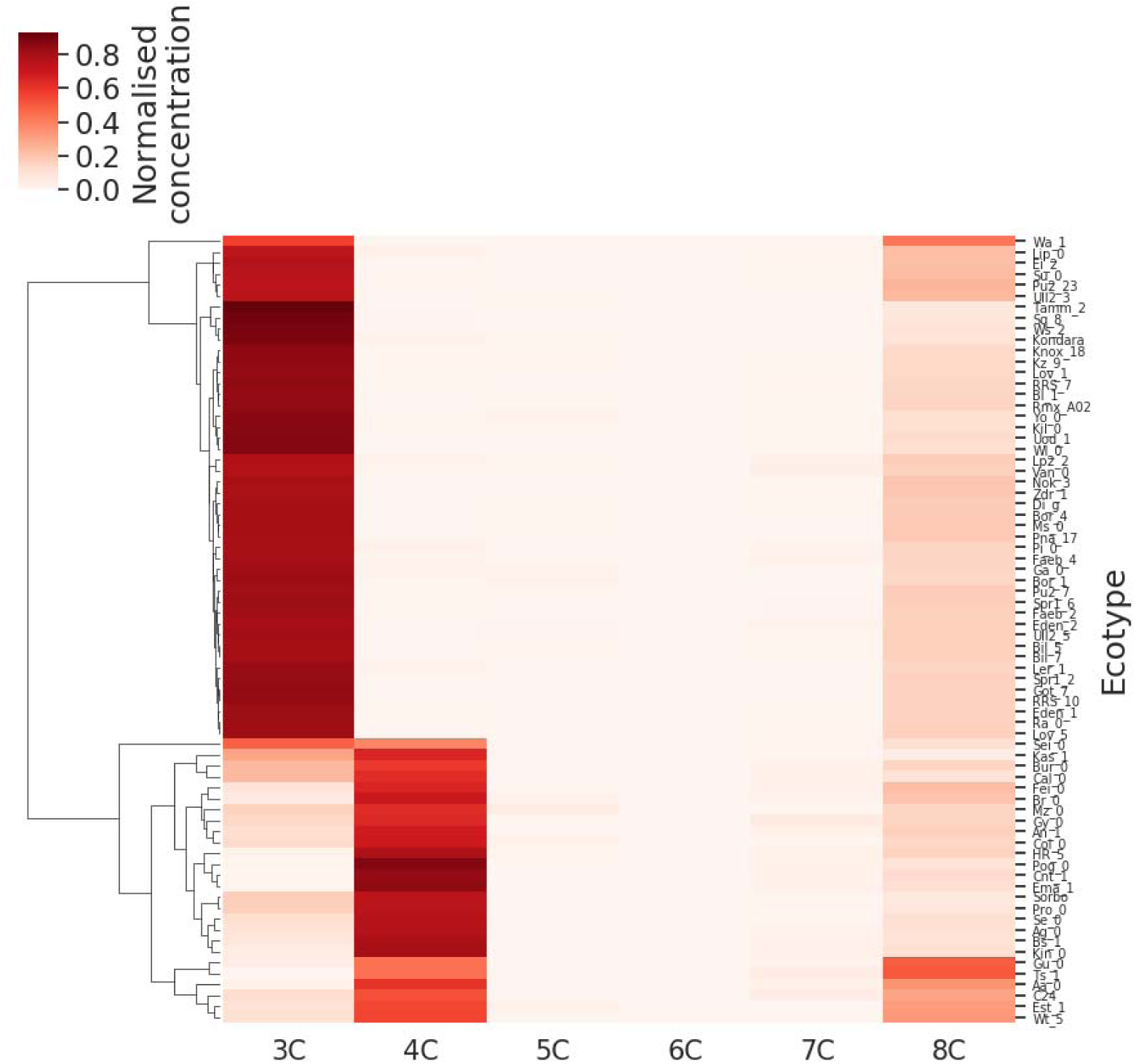
Clustered heatmap of the GSL composition across 72 different A. thaliana ecotypes. The top cluster is composed of ecotypes having high concentration of 3C GSLs, the lower cluster corresponds to the ecotypes accumulating high concentrations of 4C GSLs.

**Figure 3:**
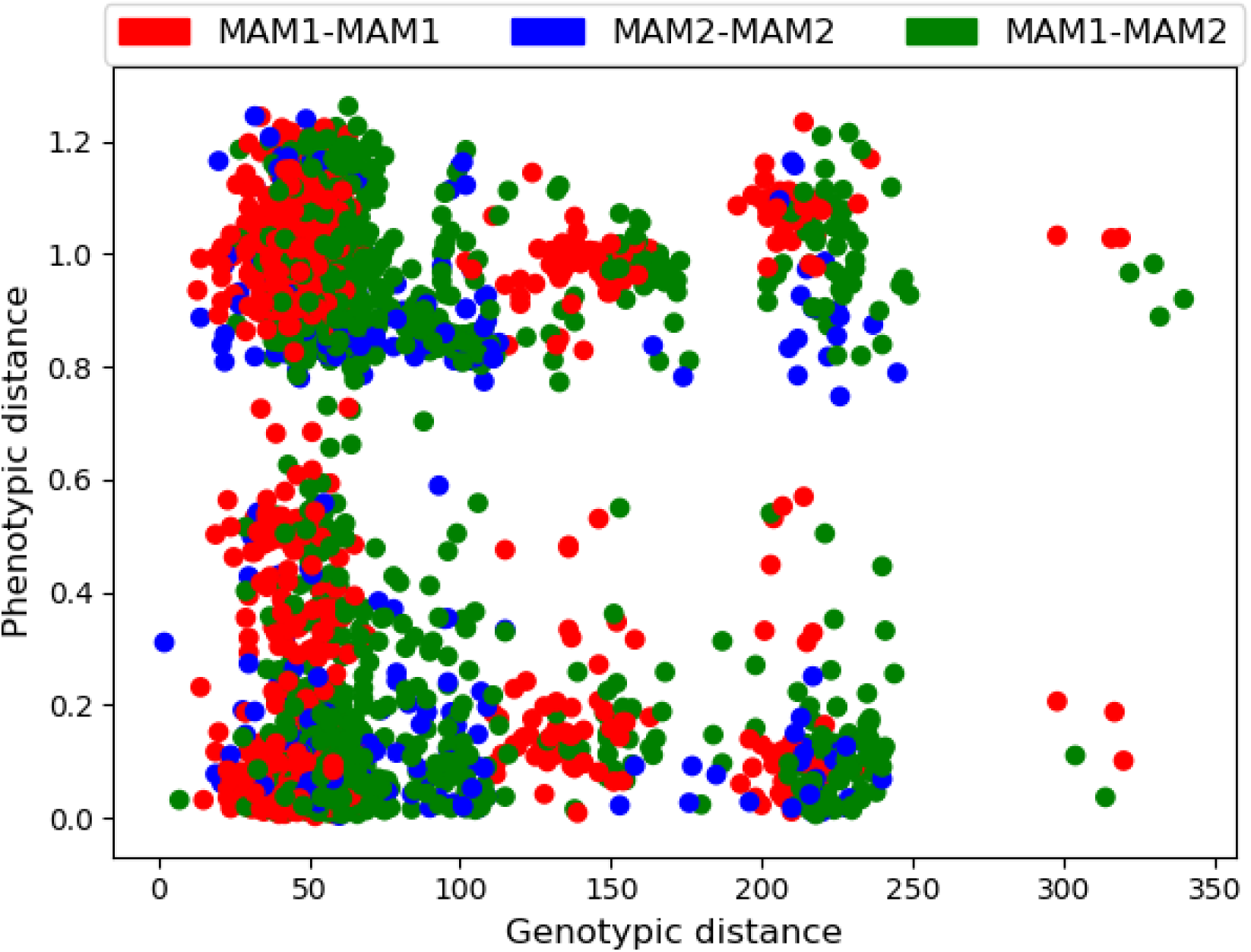
The genotypic versus phenotypic distance. Every ecotype is assigned to possess either MAM1 or MAM2, based on the sequence similarity of annotated MAM1 gene. Each dot represents a pair of ecotypes. The colours red and blue denote pairs, where both ecotypes show high similarity to the coding region of MAM1 and MAM2 sequence, respectively, the green dots denote heterogeneous pairs.

### Diversity of GSL enzyme-coding genes

To investigate the diversity of metabolic genes of GSL biosynthesis, we investigated the levels of amino acid and nucleotide polymorphism across the 72 *A. thaliana* ecotypes by calculating the average Shannon entropy (Shannon) *H* across the gene length (Figure 4A and B). The analysis revealed that some of these enzymes are highly diverse while others remain conserved across different ecotypes. Interestingly, the diversity seems to be independent of steps of GSL biosynthesis in which the enzymes are active. From the diversity of amino acid sequences (Figure 4A), *FMO-GSOX1* enzyme exhibits the highest diversity (entropy), while the lowest diversity is found in *SOT17*. Among the enzymes active in the chain-elongation pathway, *MAM1* shows the highest diversity while *BAT5* shows the low diversity. However, a further analysis of the diversity in the nucleotide sequences of the metabolic genotypes showed a high level of polymorphism in BAT5 (cf. Figure 4B), which was not reflected in the diversity of amino acid sequences. Indeed, most genes show only a slightly lower diversity in amino acid variation than nucleotide variation, which reflects the degeneration of the genetic code (Figure 4C). A plausible explanation for the low amino acid variation in BAT5 could be the specificity towards a variety of chain-elongated substrates of GSL biosynthesis (Halkier & Gershenzon, 2006). The low diversity of BAT5 could be linked to its function in transport of a diverse range of compounds that are a part of aliphatic GSL biosynthesis and Met-salvage pathway (Gigolashvili et al., 2009; Sauter, Moffatt, Saechao, Hell, & Wirtz, 2013), thus, mutations in the coding region of BAT5 may impair the functioning of both pathways. In contrast to *BAT5*, *FMO-GSOX1* shows a low diversity in the nucleotide sequences of 72 genotypes but reflect a high diversity in the amino acid sequences. This is a clear example of preferential accumulation of non-synonymous mutations, which alter the amino acid sequence of an enzyme.

**Figure 4:**
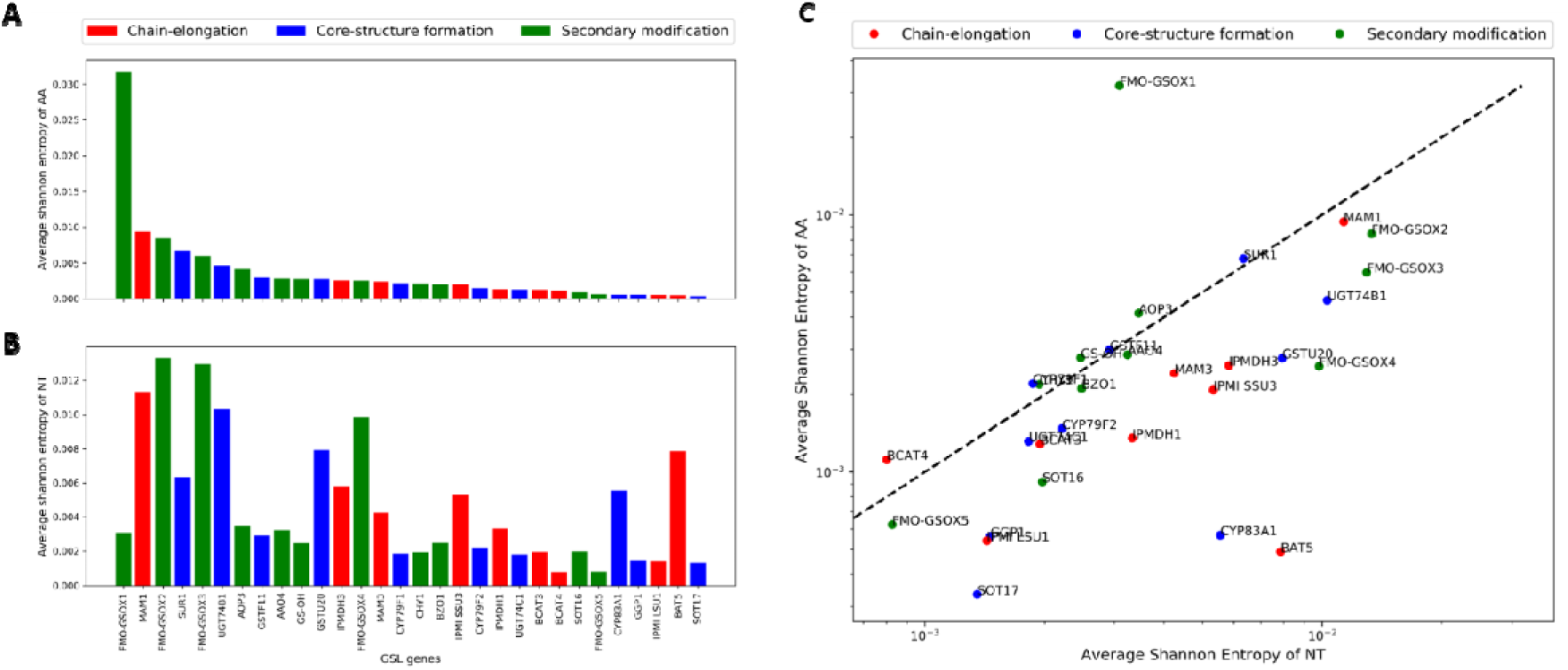
Diversity of GSL genes from 72 A. thaliana ecotypes. (A) The bars represent the diversity of the amino-acid (AA) residues of respective GSL genes. The bars are colour coded to denote genes from the chain-elongation process, core-structure formation and secondary chain modifications by red, blue and green colours, respectively. (B) The bars represent the diversity of the nucleotides (NT) of respective genes. The bars are colour coded as in (A). (C) Diversity of NT residues plotted against the diversity of AA residues of respective GSL genes. Each dot is colour coded to denote genes as in (A) and (B). The black dashed line is a linear regression line.

High diversity of *MAM1* could be a consequence of incorrect annotation of *MAM1*/*MAM2* enzymes across 72 *A. thaliana* ecotypes. Thus, we analysed the diversity of GSL enzymes across the MAM1 ecotypes (blue branch of Figure 1) and MAM2 (red branch of Figure 1) ecotypes, separately. We did see a reduction in the diversity of MAM1 and MAM2 enzymes (cf. Supplementary Figure 1 and Supplementary Figure 2). Nevertheless, *MAM1* and *MAM2* are still the most diverse enzymes of chain-elongation pathway.

### Polymorphisms in the active sites of MAM enzymes

To get a clearer understanding of the effects of the localisation of polymorphic amino acid residues in the active sites of the metabolic enzymes, we extracted the information about the active sites from the NCBI’s conserved domain database (Marchler-Bauer et al., 2015). For example, the amino acid positions 93, 94, 97, 124, 162, 164, 186, 227, 229, 231, 257, 259, 260, 261, 262, 290, 292, and 294 are known to be key for activity of MAM synthases, based on the database and a crystal structure (Kumar et al., 2019; Marchler-Bauer et al., 2015; Petersen et al., 2019). We refer to the amino acid positions from 93 to 294 as active region of the enzyme. We have found that MAM synthases exhibit a maximum of 13 and 3 polymorphic residues in the active region of MAM1 and MAM3, respectively. Figure 5(A) and (B) show pairwise comparisons of polymorphisms in the active region of MAM synthases against the genotypic distances of 72 *A. thaliana* ecotypes. Furthermore, we recorded the polymorphisms at different positions in the active region of MAM synthases (see Figure 5(C)). The amino acid residues at position 98, 99, 132, 138, 139, 147, 165, 173, 177, 187, 228, 245, 257, 258, 271, 289 and 290 of the *MAM1* and positions 156, 231, 241 and 242 of *MAM3* accumulate polymorphic residues across the 72 Arabidopsis ecotypes. Polymorphisms in the active sites of an enzyme, in principle, can change the catalytic properties of the enzyme (Kroymann et al., 2001; Kumar et al., 2019; Petersen et al., 2019). However, the quantitative effect on the enzymatic properties cannot be explained due to unavailability of enzyme abundances in these 72 ecotypes.

**Figure 5:**
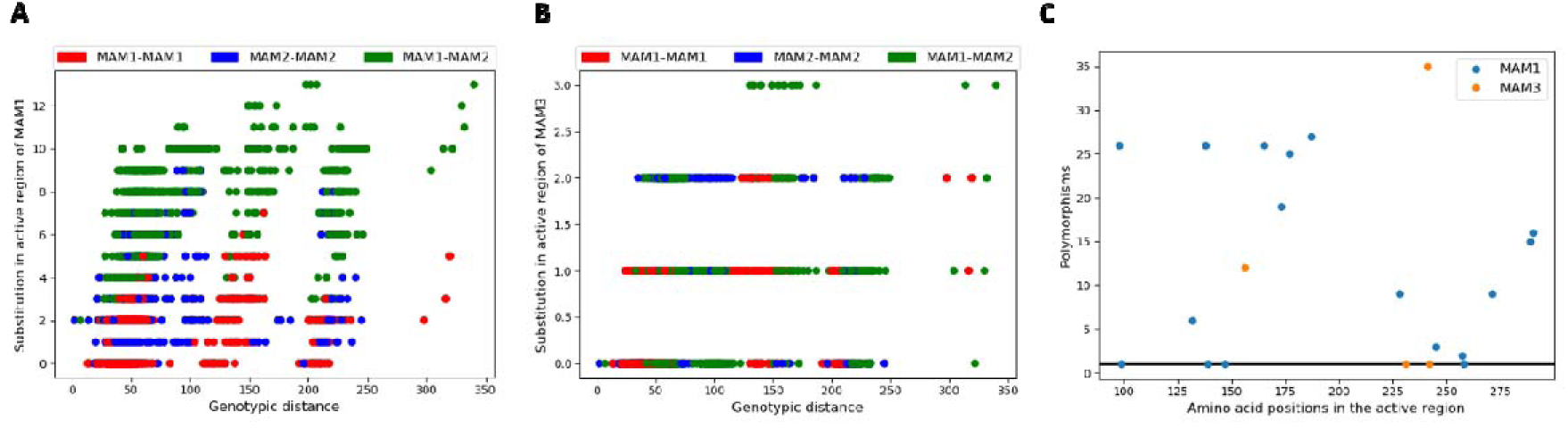
Polymorphisms in the active region of MAM synthases. (A) Substitutions in the active sites of MAM1 versus the Genotypic distances. (B) Substitutions in the active sites of MAM1 versus the Genotypic distances. (C) Polymorphisms in the active region of MAM1 and MAM3.

## Discussion

### Plasticity of the metabolic genotype and the associated GSL profiles

Glucosinolate metabolism results in a highly variable composition of individual metabolites in Arabidopsis accessions, which is reflected by a corresponding high diversity at the causal genetic level. Thus, it serves as a suitable model system to investigate the broader aspects of genotype-phenotype relationships. Allelic composition at several glucosinolate biosynthetic loci drive different glucosinolate profiles among *A. thaliana* ecotypes (Kliebenstein, Kroymann, et al., 2001). These variations, however, are often in the form of presence and/or expression of one or other copy of a duplicated gene, such as the *AOP2* and *AOP3* at the GS-ALK/GS-OHP locus (Kliebenstein, Lambrix, Reichelt, Gershenzon, & Mitchell-Olds, 2001), or the *MAM1*/*MAM2* at GS-ELONG (Kroymann et al., 2003), which complicates our understanding of how genetic variations lead to metabolic properties of the enzymes encoded by the respective genes. The Met-derived aliphatic GSLs are the most abundant form of glucosinolates in *A. thaliana* and many *Brassicaceae* crops (Agerbirk & Olsen, 2012; Benderoth et al., 2006; Halkier & Gershenzon, 2006; Kliebenstein, Kroymann, et al., 2001; Kroymann et al., 2003; Textor et al., 2004; Textor, de Kraker, Hause, Gershenzon, & Tokuhisa, 2007). The chain-elongation pathway of GSL biosynthesis is crucial for generating the chain-length diversity of aliphatic GSLs and for connecting primary and specialised metabolism. Although the evolution of core features of aliphatic GSL biosynthesis in Arabidopsis has been studied (He et al., 2011; Sawada et al., 2009; Textor et al., 2004; Wittstock et al., 2004), the molecular basis for diversity of function of MAM synthases and the role of different MAM isoforms within *A. thaliana* accessions is not sufficiently understood. The chain-length distribution of different aliphatic GSLs has so far been attributed to the presence of different MAM isoforms, namely *MAM1*, *MAM2* and *MAM3* (Halkier & Gershenzon, 2006; Kroymann et al., 2003; Textor et al., 2004, 2007). By investigating the differences in sequences of MAM synthases across *A. thaliana* ecotypes, we show that the ecotypes can be broadly classified in two groups based on the similarity to either of *MAM1* and *MAM2* genes as also expected from the genomic composition of the *GS_ELONG* locus (cf. Figure 1). Correspondingly, the GSL profiles from different ecotypes can be broadly classified into two major groups based on the phenotypic distance 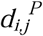 between different GSL profiles (cf. Figure 2). However, the groups classified based on either of phenotypic distances or the similarity of MAM synthases are not identical. This points to a more complicated relationship between the genotype and the associated GSL profiles. Thus, estimating the pattern of GSL accumulation based solely on the distinction between MAM1 and MAM2 enzymes is not feasible.

### Diversity of genes beyond classical QTLs

The heterogeneity in the genetic makeup of the metabolic genes across different *A. thaliana* ecotypes and their associated metabolic phenotypes are an excellent tool for investigating the mechanisms of adaptation and functional diversification (Mitchell-Olds & Schmitt, 2006; Pigliucci, 2010). Comparing the diversity of genes of GSLs synthesis we expected to find the highest diversity in genes of the chain-elongation and secondary modification, because these steps contribute highly to the diversity of the GSLs. However, surprisingly, we found that the level of diversity appears to be unrelated with the functional role of the gene within the GSL metabolic pathway (cf. Figure 4). While as expected, the least diverse enzyme was SOT17, part of the core synthesis, another enzyme of the core pathway, SUR1, was the fourth most diverse from 30 enzymes (Figure 4). This is surprising, since loss of enzymes of the core synthesis, such as *UGT74B1* or *SUR1* has a much higher impact on the total GSLs than loss of enzymes of the side chain modification (Douglas Grubb et al., 2004; Keurentjes et al., 2006; Mikkelsen, Naur, & Halkier, 2004). Thus, it seems that enzymes of all parts of GSL biosynthesis can contribute to the diversity of the metabolites. Surprisingly, while investigating the second least diverse enzyme *BAT5*, part of chain-elongation of aliphatic GSLs, we found that it is highly diverse in the nucleotide sequence (see Figure 4B). It can be a consequence of purifying natural selection that prevents the change of an amino acid residue at a given position in a multiple alignment, thus favouring an excess of synonymous versus non-synonymous substitutions. It is much more difficult, however, to explain the results of *FMO-GSOX1*, which shows a large variation in amino acid sequence derived from a relatively low variation in DNA sequence. Nevertheless, it is evident that *FMO-GSOX1* favours non-synonymous substitutions versus the synonymous substitutions, possibly linked to the function in the secondary modification of glucosinolates, responsible for large part of the structural variation. Multiple genetic analyses revealed that, in general, few key QTLs shape the metabolic phenotype. In contrast, our analysis detected diversity of the metabolic genes across the whole pathway, irrespective of their association to a major QTL. However, how far this variation in gene/protein sequence contributes to the phenotypic variation remains to be elucidated.

### The genotype-phenotype relationship

Nowhere is the contribution of subtle sequence diversity to variation in GSLs more apparent, than in the MAM genes. In *A. thaliana*, the enzyme isoforms *MAM1* and *MAM2* catalyse the formation of short-chain (*3C* and *4C*) aliphatic GSLs, whereas the isoform MAM3 catalyses the formation of both short-chain and long-chain (*5C*-*8C*) GSLs (Halkier2006). Indeed, orthologues of *MAM1* and *MAM2* are also responsible for diversity of aliphatic GSLs across a range of *Brassica* species (Kumar et al., 2019). The distinct function of the two genes was confirmed by complementation of Arabidopsis *mam1* mutant, when *MAM1* from *B. juncea* restored wild type GSL profile but *MAM2* did not (Kumar et al., 2019). Our investigation of genotypic and phenotypic distances between different Arabidopsis ecotypes showed that some ecotypes have identical metabolic genotype but exhibit high diversity in their associated metabolic phenotype, and *vice-versa* (cf. Figure 3). Therefore, the relationship between the metabolic genotypes and the phenotypes are much more complicated than a link to one of the *MAM1*/*MAM2* gene pair. Indeed, the role of individual amino acid alterations between these two genes demonstrates clearly that also SNPs can have a great effect on the phenotypes. Thus, mutagenesis of serine to phenylalanine at position 102 and alanine to threonine at 290, parts of active region of MAM1 changed the distribution of C3 to C4 GSLs (Kroymann et al., 2001) in *A. thaliana*. Alterations in four other amino acids in *B. juncea* MAM1 affected the kinetic properties of the enzyme to more MAM2-like and *vice versa* (Kumar et al., 2019). Also, in Arabidopsis MAM1 alteration of further three amino acids resulted in changes of the pattern of elongation products *in vitro* (Petersen et al., 2019). To further investigate the impact of MAM synthases on chain-length distribution of aliphatic GSLs, we analysed the polymorphisms in the active region of MAM synthases. From our analysis of the diversity of the active region of MAM synthases, we conclude that *MAM1* is highly variable across its active region and accumulate up to 13 polymorphic amino-acid residues at 17 different locations in the active region. Whereas, the active region of *MAM3* is comparatively conserved and only accumulates a maximum of 4 polymorphic residues at 3 locations in the active region, naturally (cf. Figure 5). It is known that polymorphisms in active site of MAM synthases change the specificity of a metabolic enzyme towards respective substrates of aliphatic GSL biosynthesis (Kumar et al., 2019; Petersen et al., 2019). This results in different composition and the total accumulation of aliphatic GSLs across different *Brassicaceae* species, including *Arabidopsis thaliana* (Kroymann et al., 2001; Kumar et al., 2019; Petersen et al., 2019). The above analysis, however, only showcases one of the two possibilities by which a genotype can exhibit a metabolic phenotype. The gene regulatory networks can also change the expression of metabolic genes, which in turn changes the enzyme abundance and thus results in different metabolic phenotypes (de Kraker & Gershenzon, 2011; Kumar et al., 2019; Petersen et al., 2019). Although, numerous studies have shown that a multitude of genes and underlying regulatory processes are involved in the diversity of specialised metabolites such as glucosinolates (Chan et al., 2010; Koornneef, Alonso-Blanco, & Vreugdenhil, 2004; Kumar et al., 2019; LASKY et al., 2012; Petersen et al., 2019), interpreting the findings in the context of metabolic properties is highly challenging. This is particularly due to a missing stringent definition of the genotype– phenotype relationship, which can hardly be expected to be derivable from a single methodology but rather requires a comprehensive platform of combined experimental and theoretical strategies (DIZ, MARTÍNEZ-FERNÁNDEZ, & ROLÁN-ALVAREZ, 2012; Sharma, 2018; Weckwerth, Wenzel, & Fiehn, 2004).

## Conclusion

Altogether we show here that the control over phenotypic diversity in glucosinolates is potentially spread over the whole pathway. On the example of MAM1 and MAM2, responsible for side chain elongation of Met-derived glucosinolates, we revealed that sequence variation beyond the presence of one or the other isoform contributes to the variation in chain length. The present study thus points to the necessity to pay attention to variation beyond the classical ON/OFF features of key metabolic QTLs, for investigating the diversity of specialised metabolic pathways, such as glucosinolates. Since the recent efforts towards unravelling the genotype-phenotype relationships focus at either experimental studies with a selection of genotypes or computational approaches to correlate the observed experimental observations, it is crucial to develop frameworks that integrate multi-omics data with fundamental rules of metabolic modelling to fully understand how particular genotype is reflected in a phenotype.

## Materials and Methods

### Genotypic data

Information about the nucleotide and amino acid composition of 30 GSL biosynthesis genes from 72 *A. thaliana* ecotypes was taken from the 1001 genomes project (Jorge et al., 2016). The reason behind selecting these 72 ecotypes was the availability of Met-derived GSL composition under identical environmental conditions. To obtain the gene sequences of the ecotypes of interest, we used an inhouse R-script that converts the TAIR10 version of SNP (single nucleotide polymorphisms) files provided by the 1001 genome database into an R-object. This R-object is a sparse matrix containing the nucleotide information for each ecotype at each locus in the reference genome coded as numbers. Non-polymorphic sites are coded as 0, polymorphic ones as 1,2,3,4 or 5 depending if at the specific locus an A, C, G, T or indel was observed. From this R-Object we could extract the nucleotide sequences of a specific ecotype for each coding region of interest. To obtain the amino acid sequence we used the function *’translate’* from the R-package ‘seqinr’.

### Phenotypic data

Experimental data of Met-GSLs concentrations were obtained from Chan et al. (2010) and Kliebenstein et al. (2001). The final set of GSL data is given in Table 1. The data is composed of normalised concentrations of six aliphatic GSLs, referred to as 3C to 8C, from 72 different ecotypes of *A. thaliana* under controlled and identical experimental conditions.

### Calculation of diversity using Shannon entropy

The nucleotide/amino-acid composition of the coding regions of GSL genes from 72 *A. thaliana* ecotypes is described as a set of relative probability, *p*_*i,j*_, for the *i*^*th*^ nucleotide/amino-acid (*i* = 1, 2,…, *n*) in the *j*^*th*^ ecotype (*j* = 1,2, …,72). Then, the diversity of each position in the coding region can be quantified by Shannon entropy (Shannon, 1948),

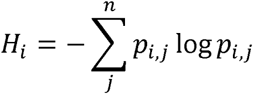

*H*_*i*_ will vary from zero, when the *i*^*th*^ nucleotide/amino-acid is same across all 72 ecotypes, to 1 when the probability is equal for observing all nucleotides/amino-acids at same locus across 72 ecotypes. Moreover, to get an estimate of diversity of a nucleotide/amino-acid sequence of length *n*, we calculate the average entropy *H*^*avg*^ as

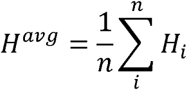

### Phylogenetic tree reconstruction

Amino acid sequences of the MAM loci from the 72 ecotypes were aligned using ‘mafft’ ver. v7.407 (Katoh & Standley, 2013) with the parameter ‘--maxiterate 1000 --globalpair -- phylipout’ to obtain a multiple sequences alignment in phylip format. This was then used as input for phyml (20120412) (Guindon et al., 2010) to reconstruct the maximum likelihood phylogenetic tree using the LG substitution model. The tree was visualised with FigTree v1.4.2 (http://tree.bio.ed.ac.uk/software/figtree/).

### Resources

The data and Python scripts used to produce the results presented in this manuscript are available with instructions at (https://gitlab.com/surajsept/GTvsPT).

## Acknowledgements

The doctoral research of SS in OE’s laboratory and research in SK’s laboratory is funded by the Deutsche Forschungsgemeinschaft (DFG) under Germany’s Excellence Strategy – EXC 2048/1 – project 390686111.

## Author contributions

SS, OE and SK planned and designed the research. SS performed the computational work and wrote the manuscript with the help of OP, OE and SK.

---

**Figure 1:**
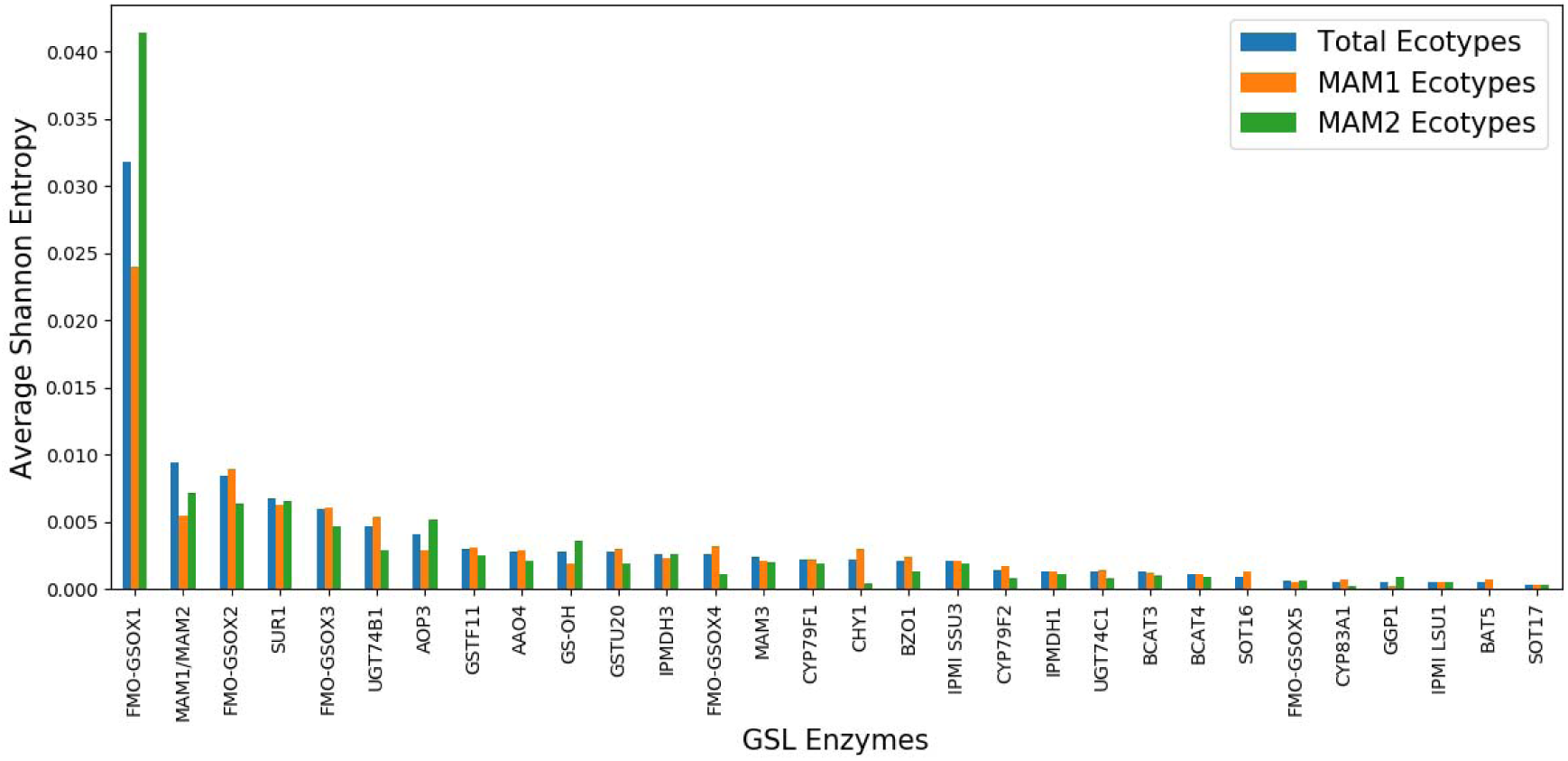
Diversity of GSL enzymes from different ecotype subgroups. The bars represent the diversity of the amino-acid residues of respective GSL genes. The blue bars denote all 72 ecotypes, while orange and green denote ecotypes having MAM1 and MAM2, respectively.

**Figure 2:**
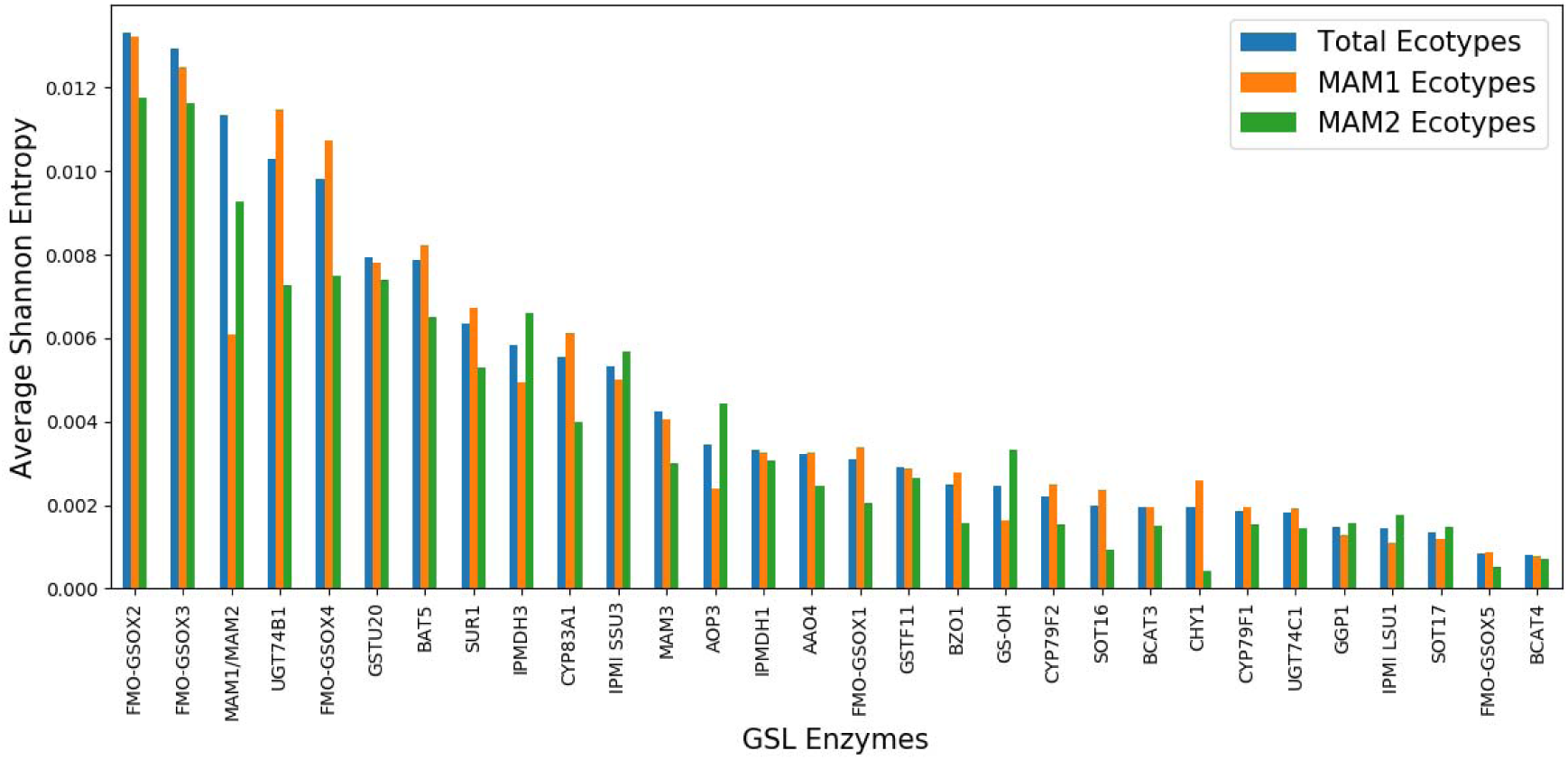
Diversity of GSL enzymes from different subgroups of ecotypes. The bars represent the diversity of nucleotide sequences of respective GSL genes. Colour coding is same as Supplementary Figure 1.

