## Supplemental Table 1 for "An integrative approach to investigate natural variation in the accumulation of aliphatic glucosinolates in *Arabidopsis thaliana*"

### Tables

Table 1: Normalised concentrations of aliphatic glucosinolates from 72 Arabidopsis thaliana ecotypes. The data is taken from (Chan et al., 2010; Kliebenstein, Kroymann, et al., 2001).

| Ecotype | 3C | 4C | 5C | 6C | 7C | 8C |
| --- | --- | --- | --- | --- | --- | --- |
| Pu2_23 | 0.741128 | 0.00652 | 0 | 0 | 0.001518 | 0.250834 |
| Bur_0 | 0.235732 | 0.583999 | 0.002074 | 0 | 0.019542 | 0.158653 |
| Gy_0 | 0.141915 | 0.6409 | 0.007476 | 0 | 0.059646 | 0.150062 |
| Gu_0 | 0.049324 | 0.437678 | 0 | 0 | 0.01782 | 0.495177 |
| Ms_0 | 0.812899 | 0 | 0 | 0 | 0 | 0.187101 |
| Knox_18 | 0.850095 | 0 | 0.008956 | 0 | 0.000757 | 0.140192 |
| An_1 | 0.116394 | 0.683634 | 0 | 0 | 0.030076 | 0.169895 |
| Lov_1 | 0.83924 | 0.009006 | 0.00278 | 0 | 0.008041 | 0.140933 |
| Se_0 | 0.122837 | 0.757981 | 0 | 0 | 0 | 0.119182 |
| Kz_9 | 0.851222 | 0.005545 | 0.006428 | 0 | 0 | 0.136805 |
| Cal_0 | 0.23823 | 0.627119 | 0 | 0 | 0.026836 | 0.107815 |
| Nok_3 | 0.798481 | 0 | 0.002419 | 0 | 0 | 0.1991 |
| Kas_1 | 0.297479 | 0.653221 | 0 | 0 | 0.005602 | 0.043697 |
| C24 | 0.125549 | 0.528313 | 0 | 0 | 0.048275 | 0.297862 |
| Ei_2 | 0.769135 | 0.005992 | 0.001686 | 0 | 0.003701 | 0.219485 |
| Mz_0 | 0.159987 | 0.628323 | 0.055078 | 0 | 0.000877 | 0.155734 |
| Kil_0 | 0.866356 | 0.012304 | 0 | 0 | 0.007637 | 0.113704 |
| Sorbo | 0.172114 | 0.744585 | 0 | 0 | 0.003106 | 0.080194 |
| Cnt_1 | 0.010444 | 0.837747 | 0 | 0 | 0.019023 | 0.132786 |
| Ag_0 | 0.10227 | 0.764187 | 0.004554 | 0 | 0.014649 | 0.11434 |
| Ts_1 | 0 | 0.447438 | 0 | 0 | 0.04507 | 0.507492 |
| Pro_0 | 0.169438 | 0.750843 | 0 | 0 | 0.005101 | 0.074617 |
| Ull2_5 | 0.81536 | 0 | 0.006728 | 0 | 0.010104 | 0.167808 |
| Bl_1 | 0.839237 | 0.001817 | 0 | 0 | 0.009083 | 0.149864 |
| Sq_8 | 0.900535 | 0.004514 | 0.002008 | 0 | 0.00646 | 0.086483 |
| Col_0 | 0.130037 | 0.6921 | 0.019605 | 0 | 0.012235 | 0.146024 |
| Fei_0 | 0.097451 | 0.65152 | 0 | 0 | 0.025647 | 0.225381 |
| Bs_1 | 0.086231 | 0.790682 | 0 | 0 | 0.023644 | 0.099444 |
| Got_7 | 0.835669 | 0.002432 | 0 | 0 | 0 | 0.161899 |
| HR_5 | 0.019516 | 0.786486 | 0.014332 | 0 | 0.019405 | 0.160261 |
| Lp2_2 | 0.77859 | 0.015955 | 0 | 0 | 0.026381 | 0.179075 |
| Ler_1 | 0.830044 | 0.015798 | 0 | 0 | 0.003286 | 0.150871 |
| Est_1 | 0.106107 | 0.5502 | 0.023744 | 0 | 0 | 0.319949 |
| Pi_0 | 0.802403 | 0.028037 | 0 | 0 | 0.014686 | 0.154873 |
| Pog_0 | 0.011563 | 0.874518 | 0 | 0 | 0.018415 | 0.095503 |
| Aa_0 | 0.018617 | 0.609043 | 0 | 0 | 0.029255 | 0.343085 |
| Ema_1 | 0.011001 | 0.84704 | 0 | 0 | 0.023573 | 0.118387 |
| Br_0 | 0.065063 | 0.701033 | 0.014885 | 0 | 0.015743 | 0.203275 |
| Ws_2 | 0.887131 | 0.005057 | 0 | 0 | 0.003123 | 0.104689 |
| Di_g | 0.806052 | 0 | 0 | 0 | 0.013755 | 0.180193 |
| Kin_0 | 0.051284 | 0.810038 | 0 | 0 | 0.022187 | 0.116492 |
| Lip_0 | 0.731449 | 0.028269 | 0 | 0 | 0.014134 | 0.226148 |
| Ra_0 | 0.825069 | 0.00355 | 0 | 0 | 0.008133 | 0.163248 |
| Wa_1 | 0.570726 | 0 | 0 | 0 | 0.006064 | 0.42321 |
| Wt_5 | 0.112922 | 0.555741 | 0 | 0 | 0 | 0.331337 |
| Bil_5 | 0.812719 | 0.004278 | 0.012447 | 0 | 0.003203 | 0.167354 |
| Faeb_2 | 0.814684 | 0.014094 | 0.002194 | 0 | 0.004514 | 0.164514 |
| Pu2_7 | 0.815383 | 0.002909 | 0.0059 | 0 | 0.000698 | 0.175109 |
| RRS_7 | 0.849395 | 0 | 0 | 0 | 0.00595 | 0.144655 |
| Su_0 | 0.748031 | 0.007874 | 0 | 0 | 0.012795 | 0.231299 |
| Rmx_A02 | 0.84428 | 0 | 0.00336 | 0 | 0 | 0.15236 |
| Ull2_3 | 0.745934 | 0 | 0.010012 | 0 | 0.001566 | 0.242488 |
| Sei_0 | 0.484848 | 0.384058 | 0 | 0 | 0.013834 | 0.11726 |
| Tamm_2 | 0.925771 | 0.004343 | 0 | 0 | 0.000435 | 0.069451 |
| RRS_10 | 0.83656 | 0 | 0 | 0 | 0 | 0.16344 |
| Uod_1 | 0.868202 | 0.001039 | 0 | 0 | 0.002911 | 0.127848 |
| Eden_2 | 0.809274 | 0.005792 | 0.007198 | 0 | 0.014818 | 0.162918 |
| Ga_0 | 0.813685 | 0.015002 | 0.021645 | 0 | 0.000556 | 0.149114 |
| Pna_17 | 0.805456 | 0 | 0 | 0 | 0.003741 | 0.190802 |
| Van_0 | 0.7742 | 0.012571 | 0.008298 | 0 | 0.039544 | 0.165387 |
| Yo_0 | 0.858786 | 0.00158 | 0.015302 | 0 | 0.003129 | 0.121203 |
| Spr1_2 | 0.831861 | 0 | 0.005072 | 0 | 0 | 0.163067 |
| Wl_0 | 0.871888 | 0 | 0 | 0 | 0.007286 | 0.120826 |
| Bor_4 | 0.805503 | 0 | 0.00845 | 0 | 0 | 0.186047 |
| Lov_5 | 0.825055 | 0.005144 | 0.000232 | 0 | 0.004937 | 0.164631 |
| Bil_7 | 0.811807 | 0 | 0.011946 | 0 | 0.007384 | 0.168863 |
| Zdr_1 | 0.788599 | 0 | 0.003087 | 0 | 0.009568 | 0.198747 |
| Faeb_4 | 0.804589 | 0.013485 | 0.003707 | 0 | 0.027481 | 0.150738 |
| Spr1_6 | 0.824565 | 0 | 0 | 0 | 0 | 0.175435 |
| Bor_1 | 0.825291 | 0.01321 | 0.016579 | 0 | 0.000617 | 0.144303 |
| Eden_1 | 0.823433 | 0.007261 | 0.00544 | 0 | 0.002107 | 0.16176 |
| Kondara | 0.882982 | 0.014853 | 0 | 0 | 0 | 0.102165 |
